# Herpes simplex virus type 1 origin binding protein UL9 tethers and loops origin- and non-origin-DNA intra- and intermolecularly

**DOI:** 10.1101/2024.10.30.621104

**Authors:** Anita F. Meier, Jan Vuckovic, Paul Girvan, Erin Cutts, Theodora Brophy, Benjamin Ambrose, David S. Rueda

## Abstract

Herpesviruses are ubiquitous human pathogens, which are the causative agent of mild to severe symptoms ranging from cold sore to nasopharyngeal carcinoma. Even though replication of the linear dsDNA genome has been studied for decades, we still lack a complete molecular understanding of its mechanism. It has been proposed, but never shown directly, that the HSV-1 origin binding protein UL9 binds two closely located binding sites within the oriS origin sequence, thereby mediating origin looping, which in turn facilitates replication initiation. Here, we used an array of single-molecule approaches to test this long-standing hypothesis directly. Surprisingly, the data show that UL9 does not loop oriS efficiently. However, we demonstrate that UL9 can form large DNA loops at non-origin sequences very efficiently, as well as tether two oriS DNA molecules intermolecularly. Contrary to the origin bending hypothesis, our findings indicate that UL9 does not loop oriS DNA, but rather may play an alternative role in replication initiation, such as tethering two separate molecules to facilitate recombination.

## Introduction

Herpes simplex virus type 1 (HSV-1) is a widespread human pathogen with a global seroprevalence of ∼65% (WHO and ^1–3^). Initial HSV-1 infection mainly occurs in the oral mucosal tissue and leads to productive replication. After an active replication phase, HSV-1 retreats into the sensory neurons of the trigeminal ganglia to establish latency^4^. While most infections remain asymptomatic, the most common symptom induced by an HSV-1 infection is cold sores. HSV-1 infections can also lead to more severe symptoms such as meningitis, keratitis, and even blindness^5^. More recent studies suggested HSV-1 as a causative agent of Alzheimer’s disease^6^. HSV-1 is one of eight human pathogenic herpesviruses within the same family and therefore shares many features with other herpesviruses such as varicella-zoster virus, Kaposi sarcoma herpesvirus, and Eppstein-Barr virus. Given the frequently severe outcomes (e.g., tumor formation) of herpesvirus infections in humans and animals, it is of great importance to understand their intricate replication mechanism.

The HSV-1 genome consists of a 152 kb linear dsDNA within the viral capsid, which circularizes upon entry into the host nucleus through an unknown recombination mechanism^7^. Genome replication is believed to occur first in an origin-dependent phase, followed by an origin-independent phase. The origin-dependent phase is usually depicted in a theta (θ) amplification mechanism, even though no direct evidence for this mechanism has been found. The origin-independent phase has been suggested to occur in a rolling-circle amplification mechanism, leading to the generation of large head-to-tail concatemers, which have been observed^8,9^. However, such concatemers could also arise from a recombination-dependent mechanism^9–11^.

The HSV-1 genome can be divided into a unique long (U_L_) and a unique short (U_S_) section, flanked by repeat regions and contains three origins of replication (**Fig. 1A**). Two of the replication origins, called oriS, are located within the repeat regions flanking U_S_, whereas the third one, called oriL, is located within the unique region of U_L_. HSV-1 encodes seven factors essential for genome replication^12,13^. These factors are the origin-binding protein UL9, the single-strand DNA-binding protein ICP8, the helicase-primase complex (UL5/UL8/UL52), and the viral polymerase and its processivity factor (UL30/UL42).

**Figure 1:**
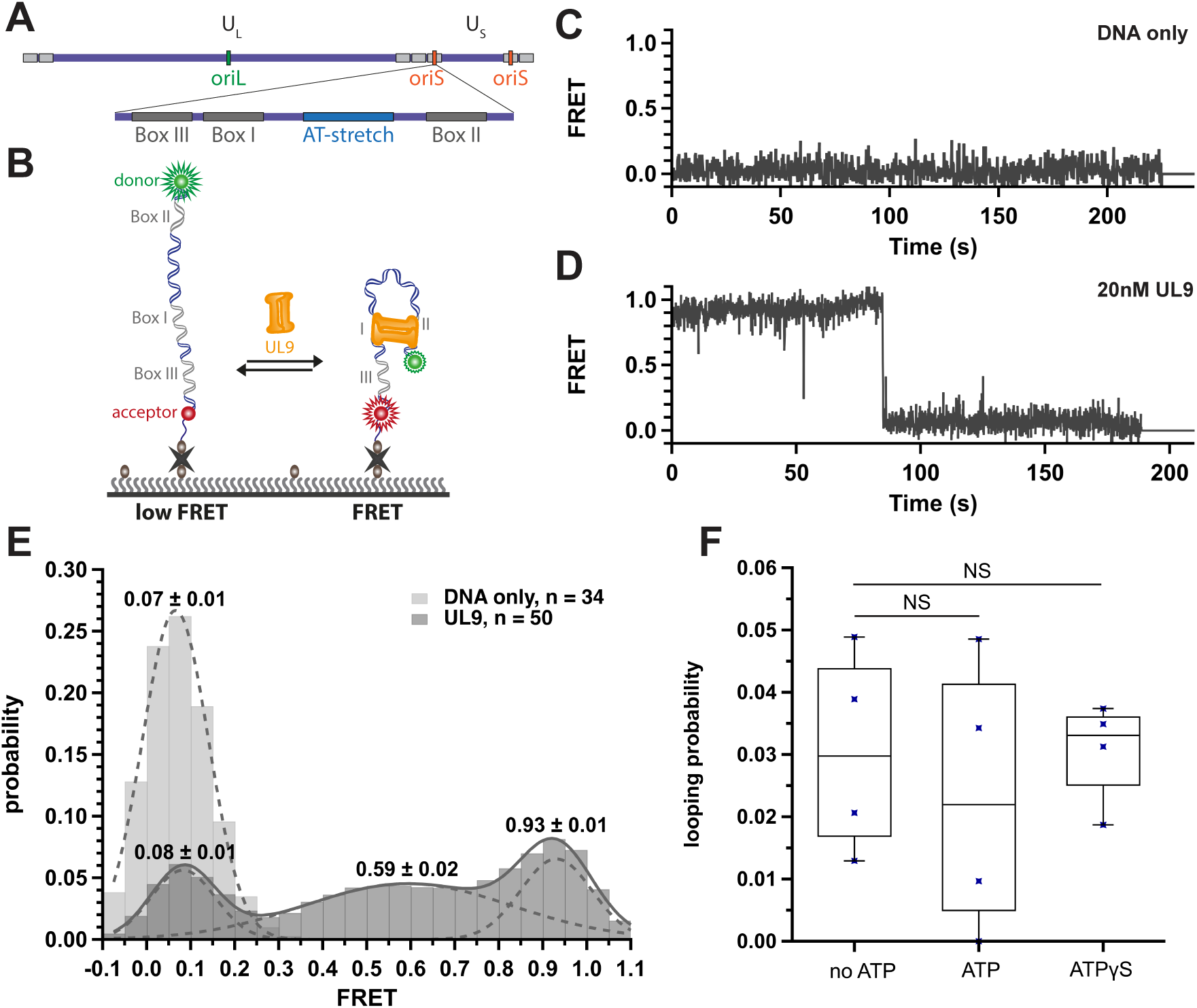
UL9 mediates origin looping at low efficiency. **A** Schematic representation of the HSV-1 genome structure including the unique long (U_L_) and unique short (U_S_) sections as well as the flanking repeat regions (grey) and the origin sequences (ori_S_ and ori_L_). The different UL9 binding sites within oriS (BoxI, BoxII and BoxIII) are indicated. **B** Depiction of the smFRET looping assay including a donor and acceptor labelled template DNA tethered to the microscope slide through biotin-neutravidin interaction. **C** Example of an oriS-DNA only control FRET. **D** Example FRET trace of an oriS-DNA in the presence of 20 nM UL9. **E** Histogram of FRET efficiency only including traces showing FRET in the absence or presence of 20 nM UL9. The data was fit to gamma distributions (dashed lines) with the centre of the gamma distribution and the overall fit (solid line) indicated. **F** Boxplot of fractions of traces showing FRET divided by the total number of traces in the presence of 20 nM UL9 only (n = 892) and with 1 mM ATP (n = 523, p = 0.62) or ATPγS (n = 706, p = 0.98). Dots represent data from independent experiments. Two sample t-test was performed with NS for p ≥ 0.05.

How the superfamily II helicase UL9 initiates the HSV-1 genome replication remains largely unknown. The oriS origin contains two high-affinity UL9 binding sites, BoxI and BoxII, that are separated by an 18 bp AT-rich sequence (**Fig. 1A**). Adjacent to BoxI, oriS also contains an additional low-affinity binding site, called BoxIII (**Fig. 1A**). UL9 has been proposed to bind to the two oriS high-affinity binding sites (BoxI and BoxII), thereby looping and distorting the intervening AT-rich DNA structure^14^. In turn, this would facilitate access to other replication initiation factors and replisome assembly.

However, this long-standing “looping” model has never been tested directly. To test this, we turned to single-molecule approaches, which eliminate ensemble averaging, thus enabling us to image conformational dynamics in real time^15,16^. The data show that UL9 can induce oriS bending but very inefficiently. However, UL9 can induce loop formation very efficiently on large DNA templates independent of the origin sequence. UL9 has only a weak preference for the origin sequence at low stretching forces and binds stably to other sequences at all forces. Furthermore, we show that UL9 tethers dsDNA inter-molecularly, suggesting a role during recombination.

## Results

### UL9 mediates oriS bending very inefficiently

To test the looping hypothesis, we surface-immobilized a 72 bp double-stranded DNA (dsDNA) template containing the oriS sequence with all three UL9 binding sites (BoxI, BoxII, and BoxIII, **Fig. 1A**) to a passivated microscope slide (**Fig. 1B**). The oriS sequence is flanked by a FRET donor (Cy3) and acceptor (Cy5) (see Methods) to measure the end-to-end distance of the DNA. We expect a long distance (low FRET) for the DNA alone and a short distance (mid to high FRET) upon loop formation. In the absence of UL9, we only observe trajectories at FRET ∼0.07, confirming that the DNA is extended (**Fig. 1C and E**). Addition of UL9 results in no apparent FRET increases for ∼97% of observed molecules (N = 892). Only a small fraction (∼3%, **Fig. 1F**) exhibits FRET increases between 0.3 and 0.9 (**Fig.1 D, E and F**), consistent with UL9-induced DNA looping. Addition of ATP, or non-hydrolyzable ATP analog (ATPγS), did not increase the fraction of molecules exhibiting mid to low FRET (∼3%, **Fig. 1F**).

To confirm that UL9 binds the oriS DNA, we visualized UL9 binding by labeling its C-terminus with an Atto647N (a FRET acceptor) fluorophore (UL9-atto647N) through a ybbR-tag^17,18^, and the surface-immobilized DNA with Cy3 only (FRET donor)(**Fig 2A**). Using Alternating Laser Excitation (ALEX)^19^, we can simultaneously monitor UL9 binding by direct acceptor emission (**Fig. 2B and C**, red) and approximate binding location by FRET (**Fig. 2B and C**, grey). The fluorescence trajectories show that the majority of molecules (79 ± 9%, n = 316) bind UL9 readily, even in the absence of ATP. We frequently observed two photobleaching steps (**Fig. 2B and C**, red), indicating that UL9 binds the oriS template as a dimer, as expected^20,21^. Based on the dwell times in the bound and unbound states, we quantify the binding kinetics of UL9 to the oriS DNA. The resulting dwell time histograms (**Fig. S2**) reveal biphasic dissociation kinetics with a short (τ_on,1_ = 1.0 ± 0.1 s) and a long dissociation time (τ_on,2_ = 9.7 ± 0.4 s) (**Fig. S2A**). These two bound times likely correspond to two binding modes. One possibility is that UL9 dissociates from one of the Box sequences (slow, τ_on,2_) or the non-specific intervening DNA sequences (fast, τ_on,1_). Alternatively, UL9 could bind in two different orientations or conformations with different binding stabilities. The unbound dwell time histogram also shows biphasic binding kinetics with a short (τ_off,1_ = 3.0 ± 0.2 s) and a long (τ_off,2_ = 25.6 ± 5.0 s) binding time, consistent with the idea that UL9 can bind in two different modes. Addition of non-hydrolysable ATPγS decreases the average bound time (τ_on,avg_) by approximately two-fold each while the average unbound time (τ_off,avg_) remains largely unchanged (**Fig. 2D**), indicating that ATP binding does not induce a conformational change that destabilizes the bound complex. Interestingly, the FRET trajectories (**Fig. 2B and C**, grey) only show mid-to low-FRET values (<0.5) after the photobleaching of one of the UL9 acceptors.

**Figure 2:**
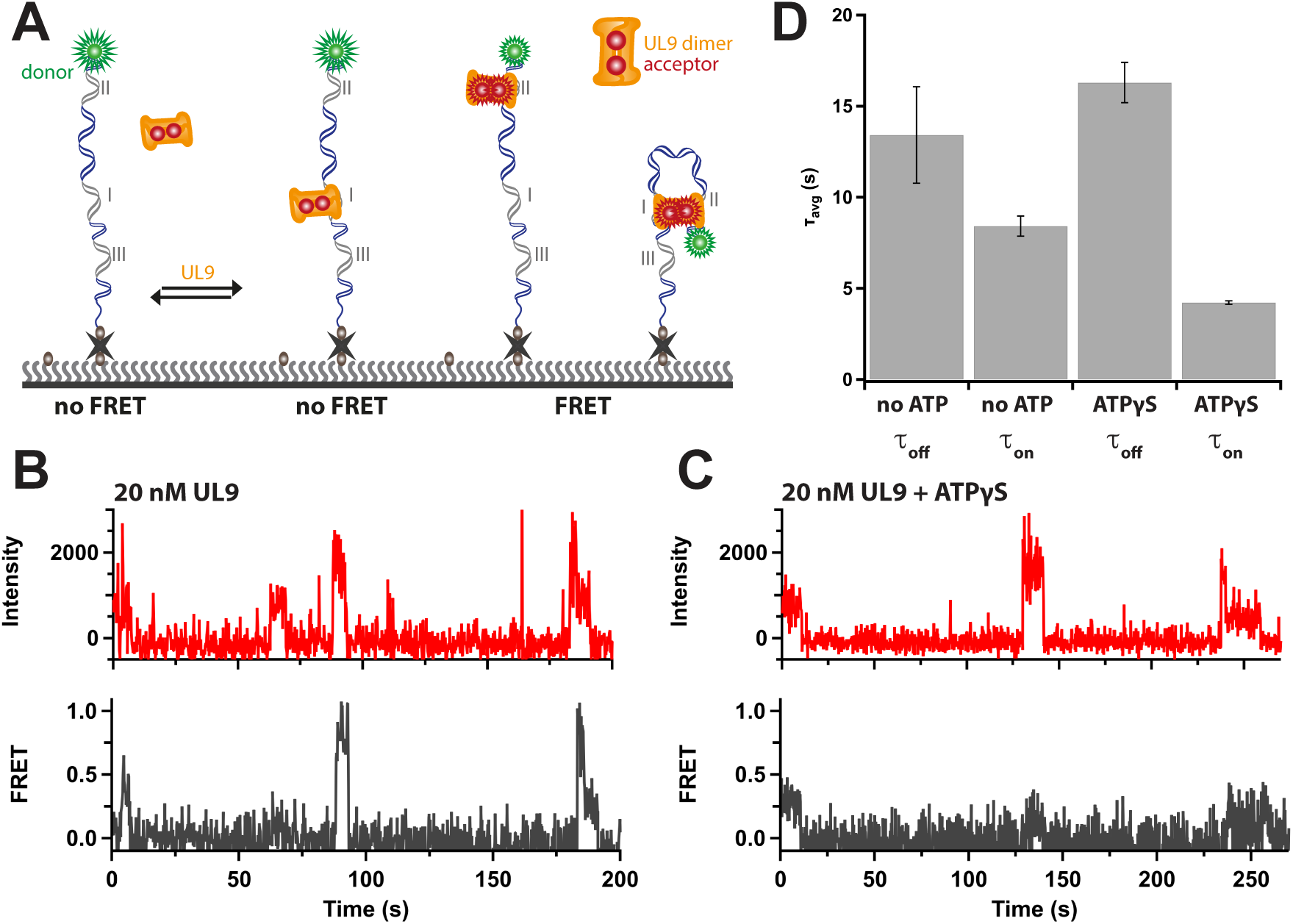
Interaction of UL9-atto647N with oriS-DNA. **A** Depiction of the experimental setup. Example traces under acceptor (red, top row) or donor (FRET, grey, bottom row) in the presence of 20 nM UL9 in the absence (**B**) or presence (**C**) of 1 mM ATPγS. **D** Bar plot of average dwell times (t_avg_) of UL9-atto647N on the oriS template at a concentration of 20 nM in the absence (unbound (t_off_, n = 1342), bound (t_on_, n = 2404)), or presence of 1 mM ATPγS (unbound (t_off_, n = 1712), bound (t_on_, n = 1918)).

Overall, our results indicate that UL9 rarely loops the oriS DNA, in agreement with the initial smFRET experiments (**Fig. 1**), raising the interesting possibility that UL9 cannot effectively bend DNAs shorter than the persistence length (∼150 bp) and may have a different function in HSV-1 replication initiation.

### UL9 efficiently induces oriS-independent loops on long DNA

To test this hypothesis, we turned to optical tweezer experiments, which enable us to observe looping on long DNA substrates^22,23^. To this aim, we prepared a DNA substrate (cos6, 47.6 kb, see Methods) encoding about a third of the HSV-1 genome and containing a single origin of replication (oriS). The linearized and biotinylated cos6-DNA was optically tethered between two streptavidin-coated beads (**Fig. 3A**) and incubated in the absence and presence of UL9 near zero force for 5-10 s before recording force-extension curves (FEC).

**Figure 3:**
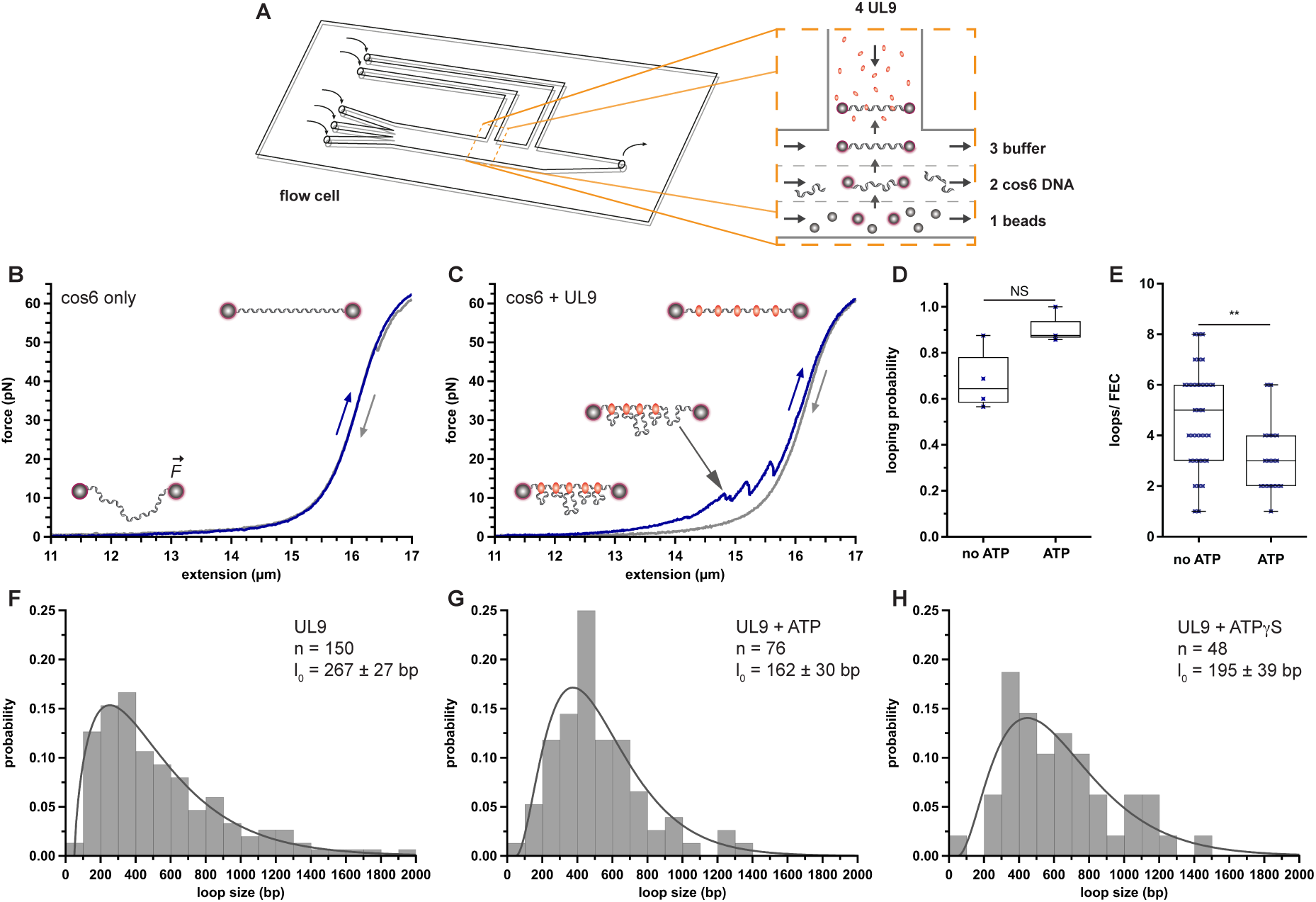
UL9 efficiently induces loops into large DNA substrates. **A** Schematic representation of the c-trap flow cell including a zoom-in of the channels. Force extension curves (FEC) of the HSV-1-derived cos6 template in buffer only (**B**) or in the presence of 10 nM UL9-atto647N (**C**) with depictions of the anticipated DNA conformation. **D** Boxplot showing fractions of rip-containing FECs in the presence of 10 nM UL9-atto647N using cos6 without (n = 52) or with 1 mM ATP (n =20, p = 0.052) as template, individual datapoints represent individual experiments from different dates. **E** Boxplot showing number of rips per FEC in the presence of 10 nM UL9 in the absence (n = 33) or presence of 1 mM ATP (n = 17, p = 0.006) as template. Two sample t-test was performed with NS for p ≥ 0.05, * for p ≤ 0.05 and ** for 0.001 ≤ p ≤ 0.01. Histogram of loop sizes in the presence of 10 nM UL9 in the absence (**F**) or presence (**G**) of 1 mM ATP or 1 mM ATPγS (**H**) using cos6 DNA.

In the absence of UL9, the cos6 DNA substrate exhibits a characteristic FEC that follows the extended worm-like chain (eWLC) model, as expected (**Fig. 3B**, blue, **and Fig. S3A**). After stretching, decreasing the bead-to-bead distance allows the DNA to refold into its original structure and follow the same FEC (**Fig. 3B**, grey). In the presence of UL9, however, we observe numerous rips in the FECs (**Fig. 3C and E and Fig. S3A**), indicating the formation of shorter contour length (L_c_) DNA likely due to UL9 looping the intervening DNA^22^, consistent with previous electron microscopy experiments^24^ (Makhov 1996). Since the cos6 DNA template contains a single origin of replication, the presence of multiple rips in a single DNA pull (on average ∼5, **Fig. 3E**) suggests that UL9 can induce oriS-independent loops within the cos6 DNA. Interestingly, looping the cos6 DNA is very efficient (∼70%, **Fig. 3D**). Next, we assessed the loop-size distribution in the absence of ATP and found that it follows the expected gamma shape^25^ with a characteristic loop length of ∼270 bp but reaching maximum loop sizes of ∼2 kbp (**Fig. 3F**). Addition of ATP, or non-hydrolyzable ATPγS, does not change the looping probability significantly (**Fig. 2D**), but decreases the average number of loops per DNA to ∼3 (**Fig. 3E**), decreases the characteristic loop length to ∼160 and 195 bp, respectively, and the maximum loop size to under 1.5 kbp (**Fig. 3G and H**). UL9-induced loops rupture forces cluster around 5 to 10 pN in the absence or presence of ATP or ATPγS (**Fig. S3B - F**), suggesting that ATP binding or hydrolysis does not stabilize oriS-independent loops. Taken together, our data show that UL9 efficiently induce large oriS-independent DNA loops, which are destabilized by ATP binding.

### UL9 shows a mild oriS sequence preference only at low force

Since UL9 appears to form oriS-independent loops, we sought to test whether UL9 has any sequence-specificity on the cos6 DNA, which contains a single oriS sequence located 7.2 kbp from the center of the linearized substrate (**Fig. 4**). To visualize UL9 binding, we used UL9-atto647N, such that binding appears as bright fluorescent dots on pre-stretched cos6 DNA at 2 pN force. The resulting kymographs (**Fig. 4A**) show that UL9 binds throughout the DNA without noticeable free diffusion^26^, revealing several oriS-independent binding events. The DNA can be trapped in two possible orientations, making it challenging to determine the oriS position on each molecule. Therefore, to examine the spatial distribution of UL9 foci on the DNA, we plotted fluorescence intensity as a function of radial distance from the mid-point of the DNA, as described^27^. At low stretching force (2pN), we see UL9 binding throughout cos6 DNA with a minor peak at the expected oriS location (7.2 kpb, **Fig. 4A**). However, this preference is lost at higher forces (>5 pN, **Fig. 4B, S3G and H**).

**Figure 4:**
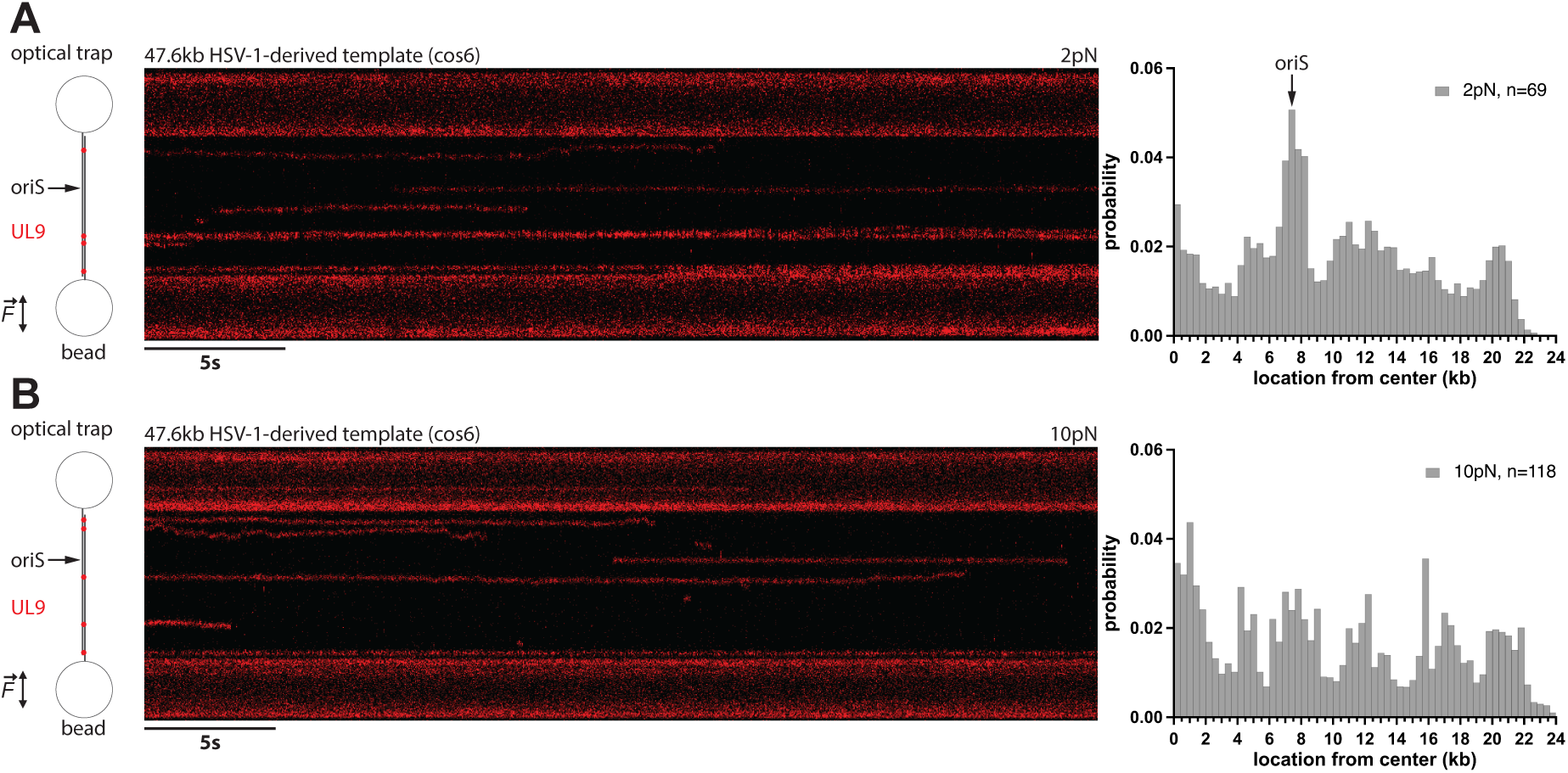
UL9 shows slight binding preference to the oriS origin sequence only at low forces. Schematic depiction of the bead-tethered cos6 DNA template held by optical tweezers and interacting UL9-atto647N (left). Example kymographs of UL9-atto647N binding (red traces) to different sites on the cos6 template DNA held at 2 pN (**A**) or 10 pN (**B**) are shown. Histograms of binding sites of UL9-atto647N to cos6 DNA plotted from the centre of the DNA held at 2 pN (**A**) or 10 pN (**B**). Location of oriS sequence is indicated (7.2 kb from the centre).

### UL9-inducued loops are static

To assess the dynamics of UL9-induced loops, we used a TIRF-based assay with surface-tethered cos6 DNA on passivated slides (**Fig. 5A**), as described^28,29^. In the absence of UL9, surface-tethered and sytox orange-stained DNA molecules show evenly distributed signal along the length of the DNA, indicating a lack of local compaction of the DNA (**Fig. 5B**). In the presence of UL9-atto647N, we observed clear accumulation of fluorescence density at various spots on the template DNA, consistent with the idea that UL9 induces local DNA compaction (**Fig. 5B and C and Fig. S4A**). Furthermore, the atto647N signal overlaps with the compacted high-intensity DNA spots (**Fig. 5B and Fig. S4A**). None of the observed high-intensity spots (DNA loops) showed migration along the DNA, and the spot intensities remained constant over time, indicating that UL9-induced loops are static (**Fig. 5C**). In one case we observed the rapid step-wise emergence of a novel loop (**Fig. S4B and C**). Together, these results suggest that UL9 forms DNA loops by rapidly tethering two distant sites on the template DNA, and remains bound statically.

**Figure 5:**
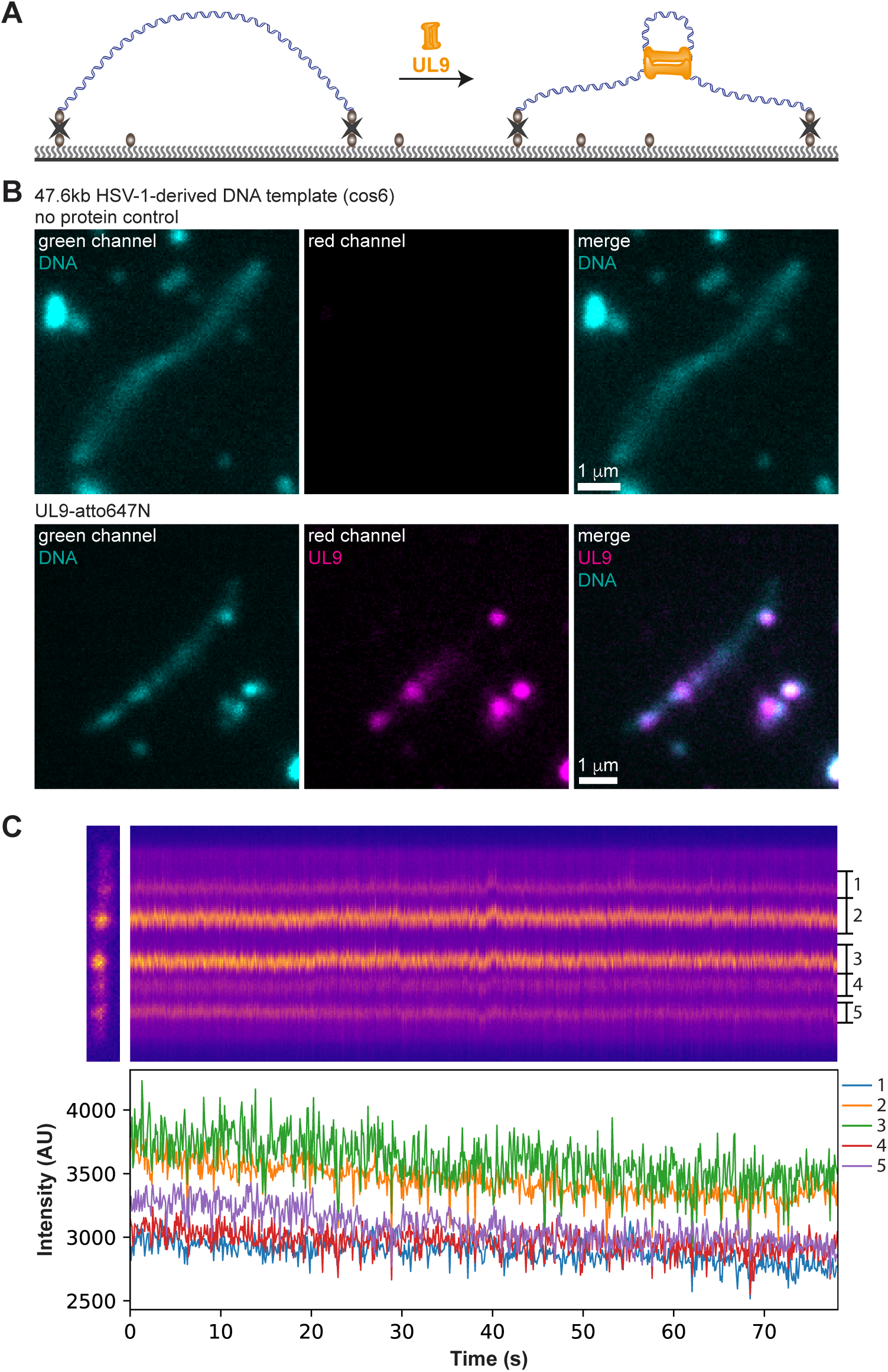
DNA loops induced by UL9 are static. **A** Schematic depiction of the experimental setup showing double-tethered template DNA. **B** Example microscopy images of sytox orange labelled cos6 DNA in the absence (top row) or absence (bottom row) of 10 nM UL9-atto647N. **C** Kymograph (top) of cos6 DNA in the presence of 20 nM UL9 with the loop-containing sections indicated on the right. Intensities of the individual loop sections are shown below. Intensity was determined by calculating the intensity per pixel within individual loop sections on the DNA over time.

### UL9 tethers dsDNA in *trans*

Given that UL9 can form DNA loops by tethering two distant binding sites, we reasoned that UL9 may also be able to tether two distinct DNA molecules in *trans*. To assess this, we turned to our TIRF set-up with a surface-immobilized Cy3-labeled oriS template (**Fig. 6A and B**), and a non-biotinylated Cy5-labelled oriS dsDNA in solution. Using ALEX, we can monitor interactions between the two DNAs by directly exciting the two dyes. In the absence of UL9, we never observed any Cy5-signal colocalizing with surface-immobilized Cy3-oriS templates, showing that the two DNAs do not interact with each other, as expected. However, in the presence of UL9, we found clear Cy5 signal colocalizing with the surface-immobilized Cy3-oriS DNA (**Fig. 6B and S5A**), showing that UL9 can readily bring together two DNA molecules in *trans*. The trajectories show that UL9 tethers the two DNAs transiently and repeatedly, reminiscent of the UL9 binding trajectories (**Fig. 2**). To quantify these *trans*-tethering kinetics, we performed dwell time analysis. In line with the UL9 binding experiments, we observed biphasic exponential decays with fast and slow bound times (**Fig. S5E and F**). The resulting average bound and unbound half-lives (τ_on,avg_ and τ_off,avg_, **Fig. 6D**) are very similar to those obtained for UL9 biding, showing that UL9 captures the Cy5-oriS DNA in solution and brings it to the surface DNA. Surprisingly, the addition of ATPγS did not affect the half-lives significantly (**Fig. 6D**). Taken together, these data show that UL9 can effectively tether two separate oriS molecules, raising the interesting possibility that UL9 may be involved in facilitating recombination.

**Figure 6:**
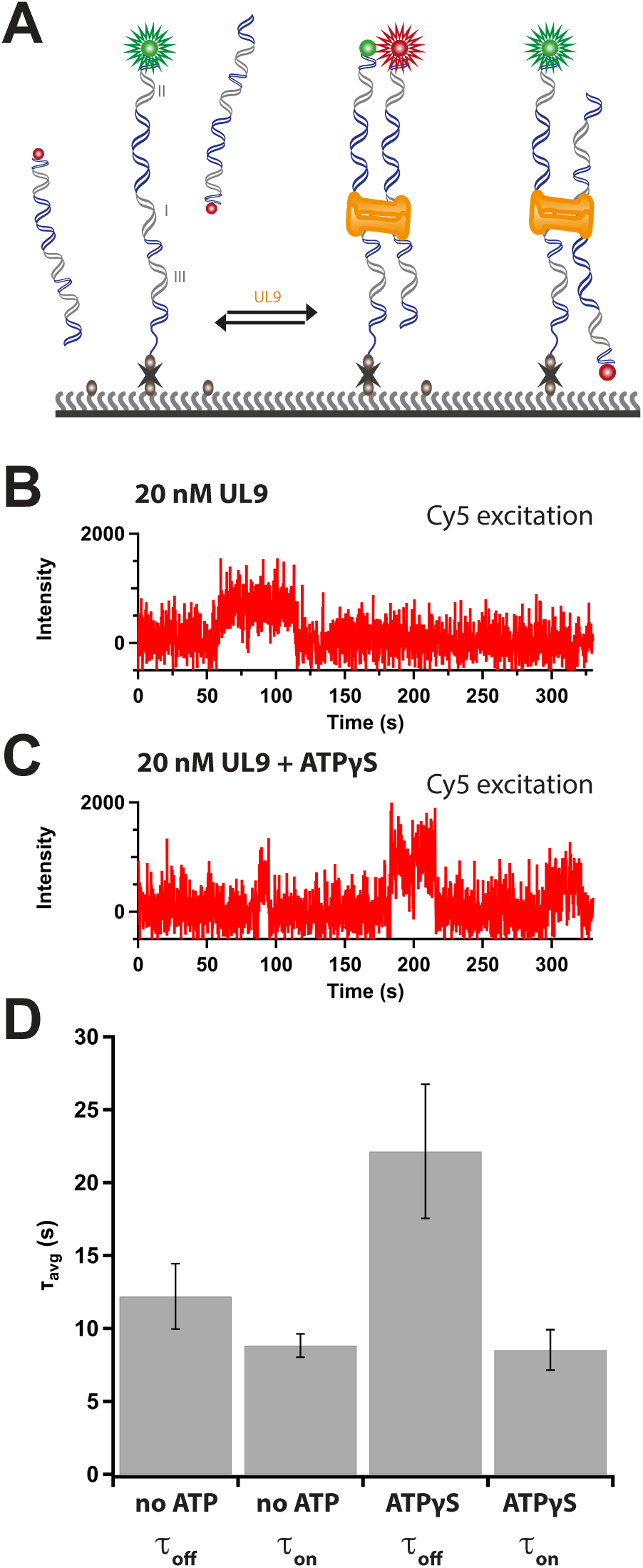
UL9 tethers dsDNA inter-molecularly. **A** Schematic of the experimental setup with donor Cy3-(donor) labelled DNA tethered to the surface and Cy5-(acceptor) labelled DNA in solution. **B** Example acceptor traces under direct acceptor excitation in the presence of 20 nM UL9 without (**C**) and with 1 mM ATPγS (**D**). Bar plot of average dwell times (t_avg_) of Cy5-DNA dwell times ± 1 mM ATPγS.

## Discussion

The molecular mechanism by which the genome of HSV-1 is replicated has not been completely elucidated. It has been postulated, but never shown directly, that the HSV-1 helicase UL9 initiates replication by looping the oriS origin of replication sequence^30,31^. Here, we have used single-molecule microscopy to test the UL9-induced oirS-looping model. Unexpectedly, the data show that UL9 does not efficiently loop a DNA containing the oriS sequence including BoxI, BoxII and BoxIII, even in the presence of ATP or non-hydrolyzable ATP analogs (Fig. 1), despite binding the DNA tightly (Fig. 2). However, using optical tweezers, we found that UL9 very efficiently induces large DNA loops independent of the origin sequence or ATP. UL9-induced formation of large dsDNA loops has previously been found by Makhov et al.^24^ by using electron microscopy but they observed that the loops were always surrounding the origin sequence. This previous finding suggested that initial UL9-mediated loop formation originated at the origin and then progressed further away from the origin. In contrast, we find that UL9-mediated loop formation is origin independent. Our TIRF-experiments suggest that UL9-induced loops are overall very static and don’t migrate. We propose that UL9 induces loops by tethering two distant locations on the DNA rather than performing a loop-extrusion mechanism. This hypothesis is further supported by our observation that UL9 is able to tether dsDNA intermolecularly. If DNA looping would arise from loop extrusion, we wouldn’t expect UL9 to be able to capture DNA from solution and tether it to the immobilized DNA. Even though it has previously been shown that UL9 is indeed able to bind to non-origin sequences^32^, we did not expect that UL9 would show stable binding to such a wide range of sequences on HSV-1-derived DNA. Furthermore, we only found a preference for the origin at low DNA stretching forces. This force-dependent sequence-specificity might be explained by the previously found interaction of UL9 with the major groove, which could be impaired by DNA distortions^33^. Based on previous work from others^31^ and the findings we present in this paper, we suggest that the origin binding protein UL9 can lead to DNA bending thereby distorting the DNA. However, this UL9-mediated oriS-bending is highly inefficient and thereby self-limiting. We hypothesize that if this UL9-mediated DNA looping occurs in the cell at the same level as we’ve observed with our smFRET assay, that this low efficiency ensures that replication is only initiated at low level during the first stage of replication thereby preventing fork collision and DNA damage resulting from fork stalling.

Based on our observation that UL9 is able to tether dsDNA intermolecularly, we propose that UL9 plays a role in HSV-1-mediated DNA recombination. Within the capsid, the 152 kb HSV-1 genome has a linear conformation. Soon after infection and entry into the nucleus, the genome undergoes end recombination and circularizes^7^. Upon replication, large concatemers can be observed, which is possibly a product of rolling circle amplification^8,9^. Interestingly, the newly replicated genomes were also found to contain inversions strongly suggesting a role of recombination during HSV-1 genome replication^9,34^. While cellular factors could be mediating recombination, previous studies suggest that recombination is mediated by viral factors such as the ssDNA binding protein ICP8 and the 5’ to 3’ exonuclease UL12^11,35–37^. UL12 was identified as a possible candidate involved in recombination based on its homology to the bacteriophage λ exonuclease Redα^38^. In bacteriophage λ, Redα and Redβ have been shown to be the only factors necessary to mediate homologous recombination (HR)^39–41^. Analogous to Redβ, the HSV-1 ssDNA binding protein ICP8 was shown to mediate ssDNA strand annealing as well as dsDNA strand invasion^42,43^. In bacteriophage λ, no helicase is required to mediate recombination. However, helicases such as HELQ in humans and Hel308 in archaea play a central role in DNA damage repair. In line with our observations for UL9, human repair proteins (e.g. Rad52^44^ and HELQ^45^) have also been found to tether DNA intermolecularly.

Interestingly, Hel308 was suggested to play a central role as regulator of homologous recombination^46^. Due to its requirement during the first phase of replication but not the second, UL9 might also take up the role of a regulator^47–49^. We hypothesize that UL9 might play a similar role to the archaeal Hel308 helicase and functions as a regulator of ICP8/ UL12-mediated recombination leading to the distinct origin-dependent and origin-independent phases of HSV-1 replication. Whether and how UL9 regulates recombination remains an open question. Supporting our hypothesis that UL9 is involved in regulating HSV-mediated DNA recombination, it was previously shown that UL9 knock-down altered the recombination pattern of adeno-associated virus genomes in an HSV-1 co-infection setup^50^. Whether UL9 enhances or suppresses recombination and what specific role the here described UL9- mediated tethering of dsDNA plays remains to be addressed in future studies.

## Acknowledgments

We would like to thank current and former members of the Rueda for insightful comments and suggestions, as well as Dr. Jaime Aspas for insightful discussions on determination of loop sizes from FECs. We also thank Bernd Vogt, Dr. Kurt Tobler and Prof. Cornel Fraefel for sharing reagents. Many thanks to Dr. Korak Ray for discussions concerning protein and DNA kinetics analysis and for advice on using tMAVEN.

## Funding

The work was funded by a core grant from the MRC Laboratory of Medical Sciences (UKRIMC-A658-5TY10, DSR), a Postdoc.Mobility fellowship of the Swiss National Science Foundation (P2ZHP3-199672, AFM) and a Marie Skłodowska-Curie Actions fellowship (101028466, AFM).

## Author contributions

AFM and DSR designed the studies. AFM conducted most of the experiments, which were analyzed by AFM and DSR. JV conducted a subset of the double-tethered DNA TIRF experiments, TB collected some of the kymographs for UL9-atto647N localisation. AFM, PG and EC prepared samples. JV, PG and BA helped generate or adapt scripts for data analysis. AFM and DSR wrote the manuscript with input from all authors.

## Competing interests

Authors declare no competing interests.

## Data and material availability

Code for single-molecule data analysis will be freely available (https://github.com/singlemoleculegroup). Correspondence and requests for materials should be addressed to AFM and DSR. All unique materials are available upon request with completion of a standard Materials Transfer Agreement.

## Notes

### Competing Interest Statement

The authors have declared no competing interest.

