## Supplementary Material for "Herpes simplex virus type 1 origin binding protein UL9 tethers and loops origin- and non-origin-DNA intra- and intermolecularly"

### Materials and Methods

#### Protein expression and purification

Sf9 insect cells were used to generate baculovirus stocks and High5 insect cells were used for protein expression. Sf9 were maintained in SF900-III SFM (ThermoFisher, 12658019) and High5 in SFX insect cell media (cytiva, SH30278) in a shaking incubator at 27°C. HSV-1 UL9 was cloned from the pCM-UL9[1] plasmid (kindly provided by C. Fraefel, UZH, Switzerland) into the pLIB vector containing a C-terminal 3C-ybbr-tev-strepII tag using Gibson cloning resulting in the pLIB-UL9-ybbR-tev-strep plasmid. In order to generate a bacmid, pLIB-UL9-ybbR-tev-strep was transformed into DH10EMBacY *E.coli* for transposition and generation of a UL9-encoding bacmid. Purified bacmid was then transfected into Sf9 insect cells to generate P0 baculovirus. After 72 hours, virus was harvested. For further amplification, the P0 UL9-baculovrius stock was passaged in SF9 cells before being used for expression in either SF9 or HighFive cells. Cell pellets were resuspended in purification buffer (20 mM HEPES pH 7.5, 250 mM NaCl, 1 mM MgCl2, 1 mM DTT, 10% glycerol) supplemented with 2 EDTA-free protease inhibitor tablets (Roche) per 50 ml and 25 U/ ml of benzonase/ nuclease (Merck) and lysed with a dounce homogeniser followed by 3 min (10 s on/ 10 s off) sonication at 20% amplitude on ice. Lysate was cleared by centrifugation followed by supernatant filtration through a 0.45 µm filter before being loaded onto a StrepTrap HP or StrepTrap XT column (cytiva), washed with purification buffer and eluted with purification buffer supplemented with 5 mM d-desthiobiotin (Merck) or 50 mM biotin (Merck) respectively. FPLC purification was performed on an Äkta pure (cytiva). Protein containing fractions were pooled, diluted 2-fold with buffer A (20 mM HEPES pH 7.5, 5% glycerol, 1 mM DTT), loaded on to HiTrap Heparin HP column (cytiva), washed with Buffer A with 200 mM NaCl, then eluted with a gradient up to 2M NaCl. UL9 was concentrated using 30K MWCO spin-concentrators (Amicon Ultra, Merck). Unlabelled protein was aliquoted and snap-frozen after the heparin column using liquid nitrogen. For labelling, concentrated (ybbR-tagged) UL9 was incubated with Sfp transferase, 5-10x excess HPLC-purified CoA-conjugated Atto647N dye, tev protease (NEB) and 10mM MgCl2 in buffer over night at 4 C or for 30min at 20°C. After centrifugation, supernatant was loaded onto a size exclusion column (Superdex 200 Increase 10/300 GL, cytiva) and fractionated with purification buffer. Protein-containing fractions were pooled and concentrated. Final protein concentrations were measured using a Qubit (Thermo Fisher Scientific) and labelling rate was assessed using absorbance (BioDrop). Labelling efficiency of UL9-atto647N was estimated to be ~10% for the initial protein batch and increased to ~50% by shortening the incubation time for protein labelling to prevent aggregation.

**Optical Tweezers**

Optical tweezer confocal microscopy experiments were performed on the commercially available Lumicks C-trap with integrated confocal microscopy and microfluidics. All channels of the microfluidics chip were first passivated with ~1 ml Pluronic F-127 (0.5% w/v in PBS) flown through at 1-2 bar. Bacterial stocks with the HSV-1 sequence containing cos6 cosmid[47] were provided by C. Fraefel (UZH, Switzerland). After extraction using a QIAGEN MiniPrep kit, the cos6 cosmid was cut with the single-cutter XbaI (NEB) leaving a 4 nt overhang on each side. The overhang was filled in using a Klenow (exo-) polymerase (NEB) in the presence of dGTP, dTTP (NEB), biotin-14-dATP and biotin-14-dCTP (Jena Bioscience) resulting in two biotins at each end[2]. Biotinylated DNA was caught in the c-trap flow cell by trapping 4.5 µm streptavidin coated polystyrene beads (Spherotech) at 0.005% w/v in laminar flow. DNA integrity was verified before each experiment by generation of force-extension curves (FECs) from 0–55 pN. For confocal imaging of atto647N, 638 nm emission wavelength and red filter 640 LP for emission was used. To record FECs in the presence of UL9, the double-tethered DNA molecule was moved into the protein channel. Then, fresh protein was flushed into the channel at low pressure (0.3 bar) for 5 to 10 s while the DNA template was held at no tension (~11 µm). FECs were recorded with 0.4 µm/ s.

### DNA oligonucleotide purification and labelling

Custom DNA oligonucleotides were synthesized by Integrated DNA Technologies (IDT) and purified by extraction from denaturing 12.5% PAGE. For labelling, the C6-dT amino-modified oligonucleotide was added to monoreactive dye (Cytiva, Q13108 (Cy3) and Q15108 (Cy5)) in 100 mM sodium bicarbonate solution and incubated over night at 4°C. On the next day, the labelled oligonucleotide was purified by high-performance liquid chromatography using an analytical C8-5 column. Purified oligonucleotides were dried using a speed-vac.

### Oligonucleotide sequences for TIRF experiments

OriS sequence included in the TIRF assays with BoxIII, BoxI and BoxII highlighted in bold in order:

5′-g**aagtgagaacg**cgaa**gcgttcgcactt**cgtcccaatatatatatattattagggcg**aagtgcgagcac**tgg

| | Description | Sequence (5' to 3') |
|---|---|---|
| **AM057** | BoxI/II/III-containing oriS template for looping and DNA capture experiments<br>Labelled with Cy3 for looping and UL9-interaction assay and labelled with Cy5 for the capture assay | /5AmMC6T/ccagtgctcgcacttcgccctaataatatat atatattgggacgaagtgcgaacgcttcgcgttctcactt c |
| **AM071** | Complementary to AM057 with a ssDNA tail of 9 adenines<br>Labelled with Cy5 for looping and unlabelled for the capture and UL9-interaction assays | /5Biosg/aaaaaaaaa/iAmMC6T/gaagtgagaacgcg aagcgttcgcacttcgtcccaatatatatatattattagg gcgaagtgcgagcactgg |
| **AM058** | Complementary to AM057 (without tail) used in the capture assay | gaagtgagaacgcgaagcgttcgcacttcgtcccaatata tatatattattagggcgaagtgcgagcactgg |

### Microscope slide passivation and flow chamber assembly

Quartz (for smFRET, UQC optics) or glass (for double-tethered DNA experiments, Menzel-Gläser, ThermoScientific) slides with custom-drilled holes and glass coverslips (24 x 50 mm, No. 1.5H, VWR) were coated with aminosilane (N-(2- Aminoethyl)-3-aminopropyltrimethoxysilane) then pegylated using methoxy-PEG-SVA (Mr = 5,000, Laysan Bio, Inc.) containing 10 % biotin-PEG-SVA (Mr = 5,000, Laysan Bio, Inc.) in 100 mM sodium bicarbonate as described previously[3]. Following passivation, slides and coverslips were stored in nitrogen at -20°C in the dark. Prior to use, slides and coverslips were equilibrated to room temperature. If necessary, tubing was glued into the holes in the microscope slides and excess tubing cut off. The flow chambers were then assembled using double-sided adhesive tape or 0.12 mm thick double sided adhesive sheets (Grace Bio-Labs SecureSeal). Flow chambers were sealed with epoxy glue.

**smFRET experiments**

Biotin-PEG-passivated home-built quartz flow chambers (see sections above for more details) were incubated with 0.1 mg/ml neutravidin in T50 (50 mM NaCl and 50 mM Tris-Cl (pH 7.5)) for 5min then blocked for 10 min with Pluronic F-127 (0.5% w/v in T50). PAGE-purified biotinylated and fluorescently labelled complementary oligonucleotides (see other sections of more details) were annealed at a concentration of 1 µM in T50 by incubation at 95°C for 45 s followed by 10-15 min annealing at room temperature and immobilised on the flow chamber at 5 pM and 2 min incubation. After DNA-immobilisation, a further biotin (1.7 µM) blocking step in UL9 buffer (50 mM NaCl, 0.1 mg/ml BSA, 40 mM Tris-Cl (pH7.5), 1mM $MgCl_2$, 0.5 mM DTT and 50 mM potassium glutamate) was performed. Imaging was performed in UL9 buffer supplemented with glucose-oxidase scavenger system (8 mg/ ml D-glucose, 1 mg/ ml oxidase and 40 µg/ ml catalase) in the presence or absence of 20 nM UL9-atto647N or 20 nM UL9 and 1 mM ATP or ATPγS. For the DNA capture assay, Cy5-labelled (non-biotinylated) DNA was provided at 10 nM in solution. Data acquisition was performed on an Olympus IX-71 microscope equipped with a homebuilt prism-TIRF module. Excitation was provided by a 532 nm or a 637 nm laser. Fluorescence was collected through a 1.2 NA, 60x water objective (Olympus) and filtered through a dual bandpass filter (FF01-577/690-25, Semrock). The fluorescence was spectrally separated using a OptoSplit II (Cairn Research) to separate donor and acceptor emission. The donor and acceptor emission were further filtered through ET585/65M and ET700/75M (Chroma) bandpass filters, respectively. The donor and acceptor images were then projected side-by-side onto an EMCCD (Andor iXon Ultra897). Data was collected as raw movies using a custom LabVIEW script. Single molecule fluorescence spots from the raw movies were localised using custom IDL scripts and converted into raw fluorescence trajectories. Raw fluorescence trajectories were corrected for bleed through of the donor fluorescence into the acceptor channel. Apparent FRET efficiencies were calculated as the ratio of acceptor intensity divided by the sum of the donor and acceptor intensities. Two mechanical shutters (LS-3, Uniblitz, Vincent Associates) were placed in the excitation path for alternating laser excitation. Frame acquisition and shutter synchronisation was obtained using a homebuilt negative edge triggered JK flip-flop circuit (SN74LS112AN, Texas Instruments) using the “Fire” output of the EMCCD as the input clock. IDL scripts were modified accordingly to locate single molecules and extract fluorescence trajectories. Trajectories were only included if they met specific criteria. For the smFRET looping experiments, to be considered in the dataset, trajectories had to include a single donor bleaching step and a minimal donor intensity of 1000 AU. Furthermore, for the experiments where the red laser was turned on at the beginning of imaging for a few frames before the green laser was turned on, the trajectories had to show a clear acceptor signal to be included in the dataset. For the UL9-atto647N binding and DNA capture experiments, where alternating laser excitation was used, the trajectories had to show clear acceptor and donor signal to be included in the dataset.

**Synthesis of λ-DNA with a single biotin at each end**

The 12 nt overhang ends of lambda phage DNA were modified to each contain a single biotin attachment based on literature[4]. 5’-phosphorylated and 3’-biotinylated DNA oligonucleotides, LDNA_biot_end_1 (5’-pAGGTCGCCGCCC-TEG-Biotin-3’) and LDNA_biot_end_2 (5’-pGGGCGGCGACCT-TEG-Biotin-3'), were purchased from Integrated DNA Technologies. The oligonucleotides (1.2 pmol) were annealed to the λ-DNA ends sequentially (0.4 pmol, Thermo Scientific SD0011) by incubating the mixture at 70 °C for 15 min in 1X T4 DNA buffer (400 µl, NEB B0202), followed by slow-cooling to room temperature. The annealed products were then ligated by adding T4 DNA ligase (320 units, NEB M0202) and 100 mM ATP (400 nmol, Thermo

Scientific R0441) and incubated at room temperature for 2 h. After annealing and ligation of both oligos, the ligase was heat-inactivated by incubating the mixture at 65 °C for 20 min.

**Double-tethered DNA experiments at TIRF**

All steps were performed on the home-built objective-based TIRF microscope with the outlet tubing connected to a syringe pump. Biotin-PEG-passivated home-built glass flow chambers (see sections above for more details) were incubated with 0.1 mg/ml neutravidin in T50 (50 mM NaCl and 50 mM Tris-Cl (pH 7.5)) for 5min then blocked for 10 min with Pluronic F-127 (0.5% w/v in T50). End-biotinylated cos6 (see c-trap section for details) or λ-DNA with a single biotin at each end was immobilized at a flow rate of ~ 5 µl/min and a total of 300 µl of ~2-4 pM DNA. Once the DNA was immobilised, slow flow rates (2-5 µl/min) were used to prevent damage to the DNA template. A second blocking step was performed using 1.7 µM biotin in UL9 buffer (50 mM NaCl, 0.1 mg/ml BSA, 40 mM Tris-Cl (pH7.5), 1mM $MgCl_2$, 0.5 mM DTT and 50 mM potassium glutamate). Immobilised DNA was imaged in UL9 buffer with 50 nM SYTOX orange (sxo, ThermoFisher) and glucose-oxidase scavenger system (8 mg/ ml D-glucose, 1 mg/ ml oxidase and 40 µg/ ml catalase) in the presence or absence of 10 nM UL9-atto647N or 20 nM UL9. To image the sxo signal, the excitation laser with the wavelength of 532 nm and to image the atto647N signal, the 637 nm wavelength laser were used. For emission, a 534/640 bandpass filter (Chroma, DC/ZET532/640m) with a compatible dichroic mirror was used. The fluorescence was spectrally separated using a MultiSplit (Cairn Research) housing the following dichroic filters: T500lpxr UF2, T635lpxr UF2 and T725lpxr UF2 (Chroma). The separated fluorescent emission was projected onto quadrants of a sCMOS (ORCA Fusion, Hammamatsu) camera. Data was collected as raw movies using HCImage Live (Hammamatsu). Imaging data was analysed using in-house python scripts. After co-localisation of the separate green and red channels using python, final images were merged, cropped and coloured using ImageJ[5]. Kymographs were generated by selecting an area covering the DNA molecule and then tiled over time. To assess the intensity of the individual loops, the area of the spots was selected and the overall signal intensity was normalised to the number of pixels. Plots were generated using home-built python scripts.

**Data analysis**

Analysis of smFRET traces for the oriS-looping assays

To generate histograms assessing the different FRET states, the sections of the traces showing FRET (or entire traces for the controls) were selected and converted into FRET efficiency traces ($E_{FRET} = I_{acc.}/ (I_{acc.} + I_{don.})$) using MATLAB (MathWorks). Histograms were generated and fitted using IgorPro8 (WaveMetrics). Transition density plots (TDP) and post synchronisation plots were generated with tMAVEN[6] using default parameters.

**Kinetic analysis of UL9-atto647N binding and DNA capture assay**

Raw atto647N/ Cy5 traces were first converted into a format that could be used by tMAVEN using an in house-matlab script. We then used python to convert the file into an .h5 file. In tMAVEN[6] we first performed a global Hidden-Markov Modelling (HMM) for all traces of each experiment (vbConsensus) assuming two states and using the default parameters. Since we only assessed binding/ no binding, we didn't distinguish any multimer binding states. After HMM modelling, we removed the traces, which were over-fitted. Dwell times from the idealized FRET trajectories were extracted using custom MATLAB (MathWorks) scripts. The data was then plotted as histograms with IgorPro8 (WaveMetrics) and fit to a single or double exponential distribution to determine k and τ (1/k).

Single exponential: $f(t) = A \exp(-kt)$

Double exponential: $f(t) = A_1 \exp(-k_1t) + A_2 \exp(-k_2t)$

Standard deviation of τ was determined as follows: $d\tau = dk/ k^2$

Average lifetime ($\tau_{avg}$) for UL9-atto647N binding or DNA capture was calculated from the double exponential fit as follows:

$T_{avg} = (A_1 \tau_1^2 + A_2 \tau_2^2) / (A_1 \tau_1 + A_2 \tau_2)$

**Calculation of loop sizes from c-trap force-extension curves (FECs)**

FEC data was extracted from .h5 files using a home-built python script[7]. Loop sizes were then assessed in MATLAB (MathWorks) using a home-built script. The PeakFinder function was used to first identify the coordinates (DNA length and rupture force) of the rips and then the corresponding point at the same force (DNA length after loop rupture). The difference between the initial length at the rupture point and the DNA length after the rupture was determined and converted into an approximate number of base pairs with 1bp = 0.338 nm[8]. Plots were generated with IgorPro8 (WaveMetrics). For the fit shown in the histogram of the loop sizes, we used the following equation (generalized gamma distribution): $f(l) = A * l^{(n-1)} * e^{(-l/l_0)}$

**Determination of UL9-atto647N binding site from c-trap kymographs**

Trace length and location from kymographs of UL9-atto647N binding to the HSV-1-derived cos6 template were analysed using a home-built python script. Distance from the centre of the template in each frame from the entire trace was then calculated in MATLAB (MathWorks) and histograms were plotted using IgorPro8 (WaveMetrics).

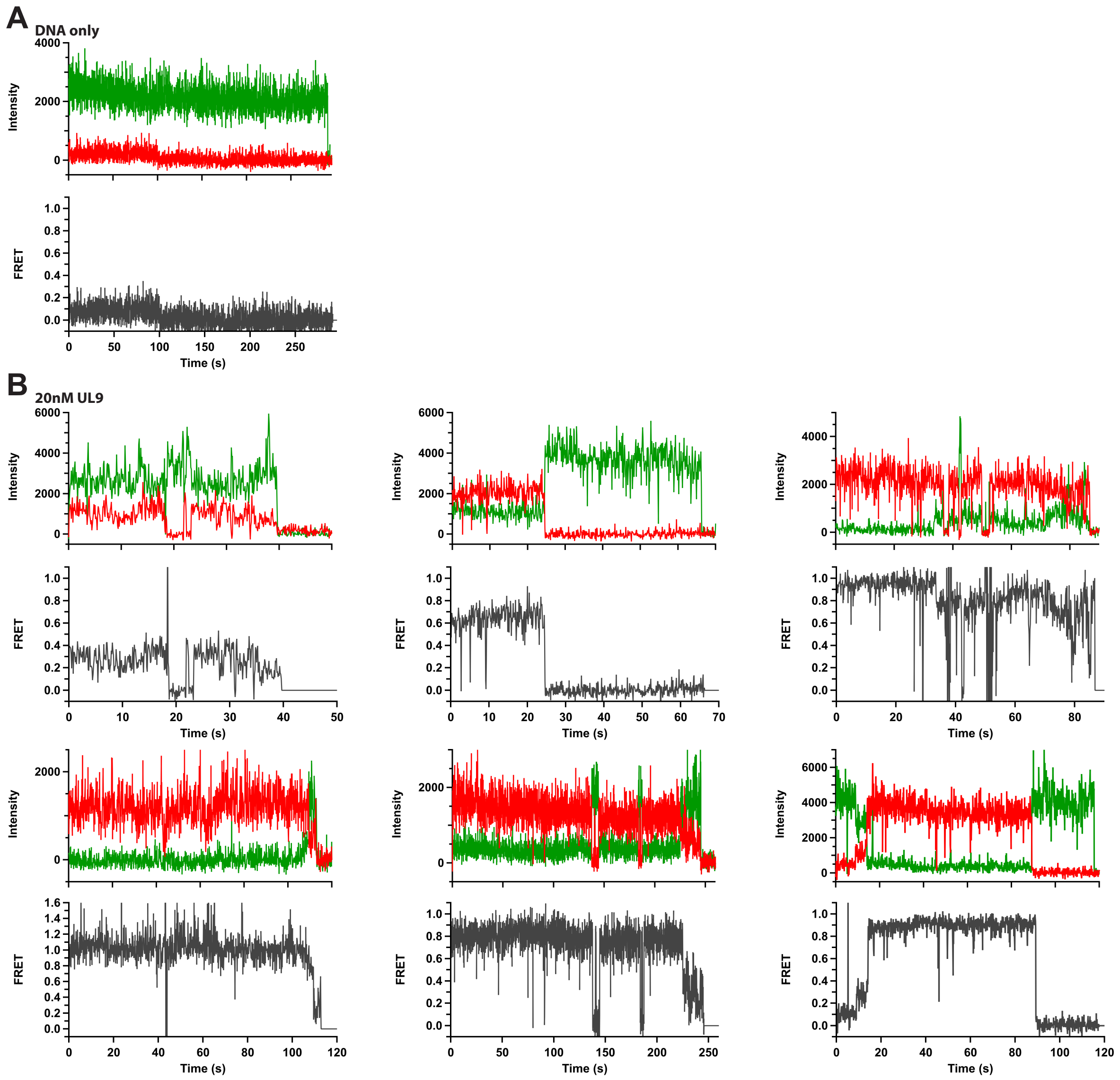


**Figure S1: Additional data of the smFRET oriS-looping assay.** Example traces of oriS DNA in buffer (**A**) or in the presence of 20 nM UL9 (**B**) with donor (green) and acceptor (red) signal in the panel on the top row and the FRET efficiency (grey) on the bottom row.

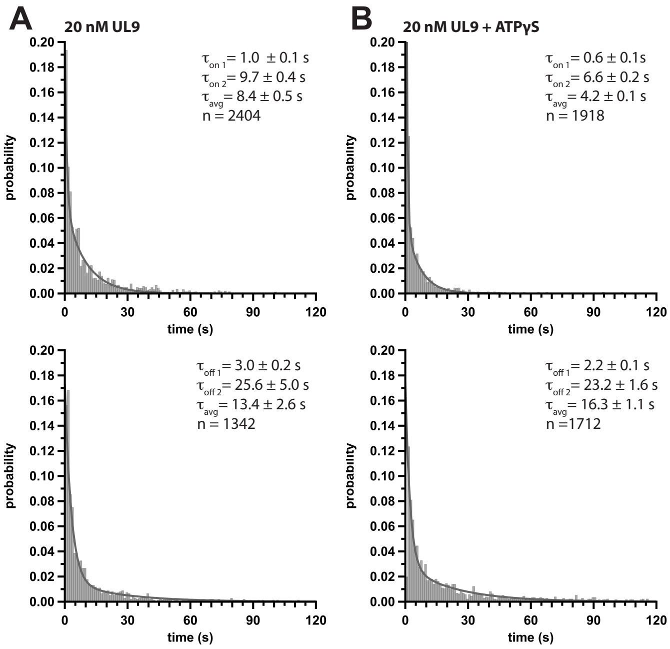


**Figure S2: Dwell time determination of UL9-atto647N on oriS.** Histograms of UL9-atto647N dwell times in the bound (top row) and unbound (bottom row) state in the absence (**A**) and presence (**B**) of 1 mM ATPγS.

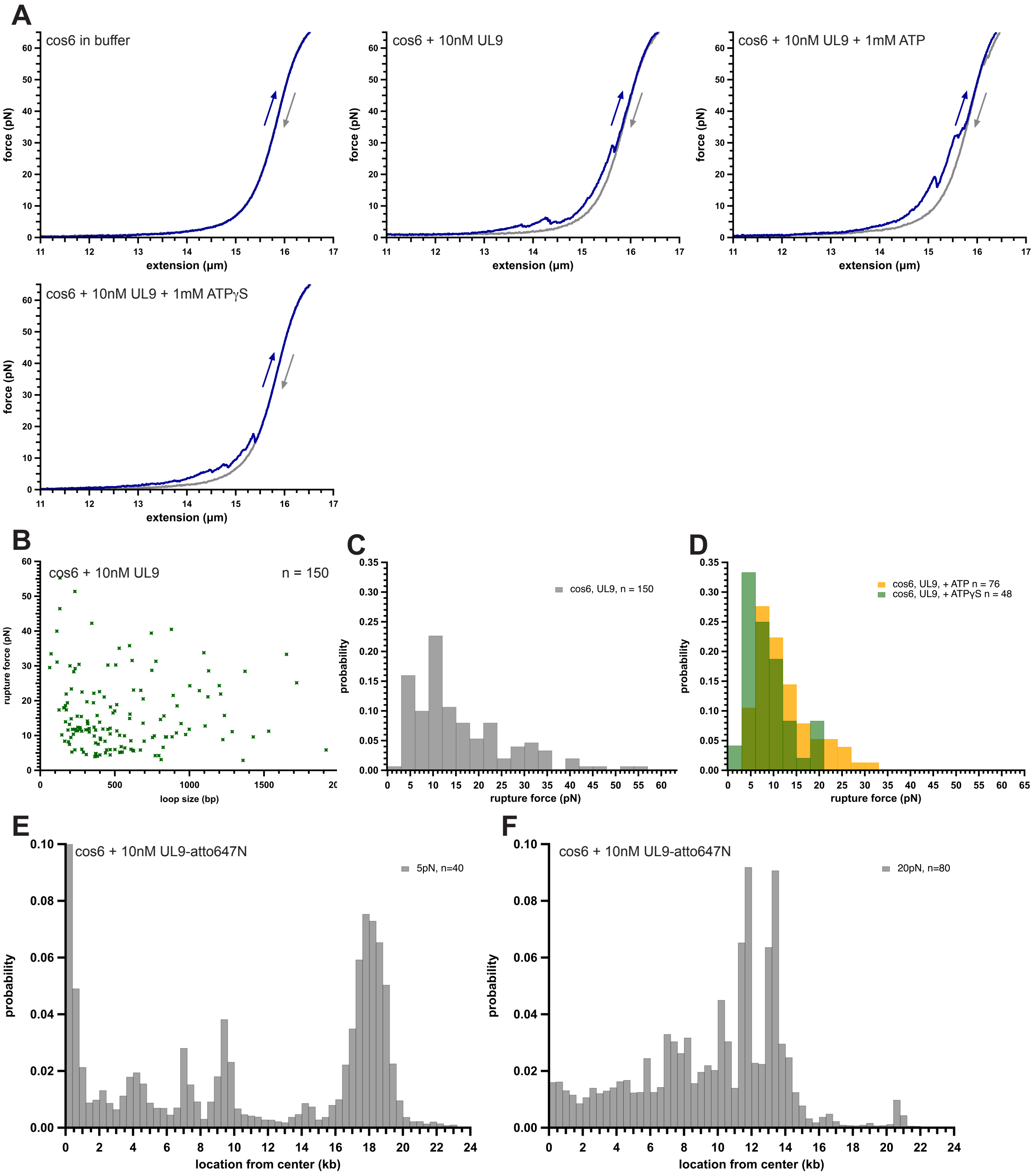


**Figure S3: Additional data on UL9-induced loop formation at the c-trap. A** Force-extension curves (FECs) of cos6 or in buffer only and in the presence of 10 nM UL9 or 1mM ATP/ ATPγS as indicated. **B** Dot plot of loop size vs. rupture force of cos6 in the presence of 10 nM UL9. Histograms of loop rupture forces in the presence of 10 nM UL9 using cos6 without ATP (**C**) and with 1 mM ATP (yellow) or ATPγS (green) (**D**). Histograms of UL9-atto647N binding location on cos6 DNA plotted from the center of the template held at 5 pN (**E**) or 20 pN (**F**). Number of traces are indicated (n).

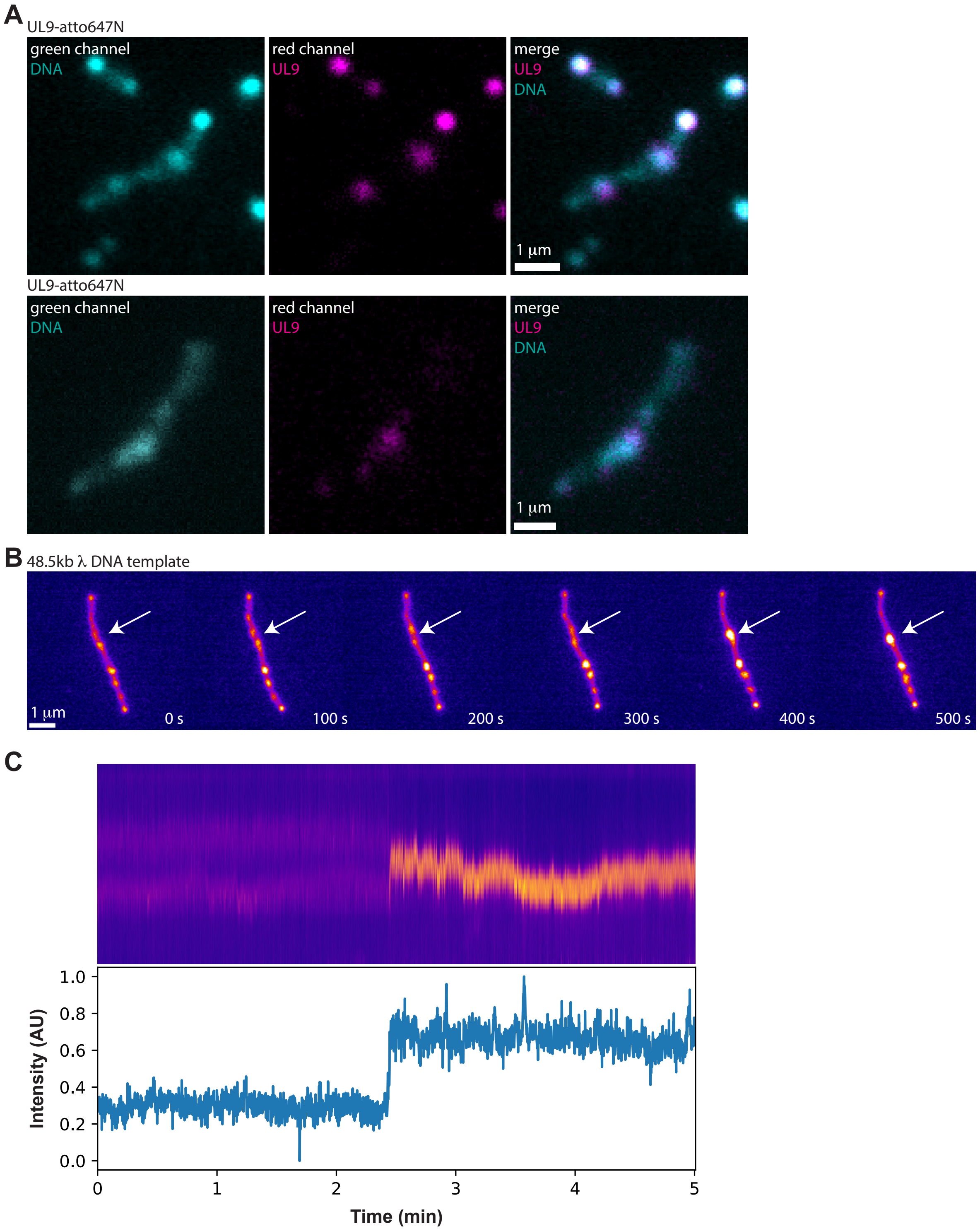


**Figure S4: UL9 induced loops double tethered DNA. A** Microscopy images of cos6 DNA in the presence of 10 nM UL9-atto647N. **B** Microscopy images at indicated timepoints of an emerging loop (arrow) on double-tethered λ DNA. **C** Kymograph of a zoomed-in region of the emerging loop with the pixel intensity below.

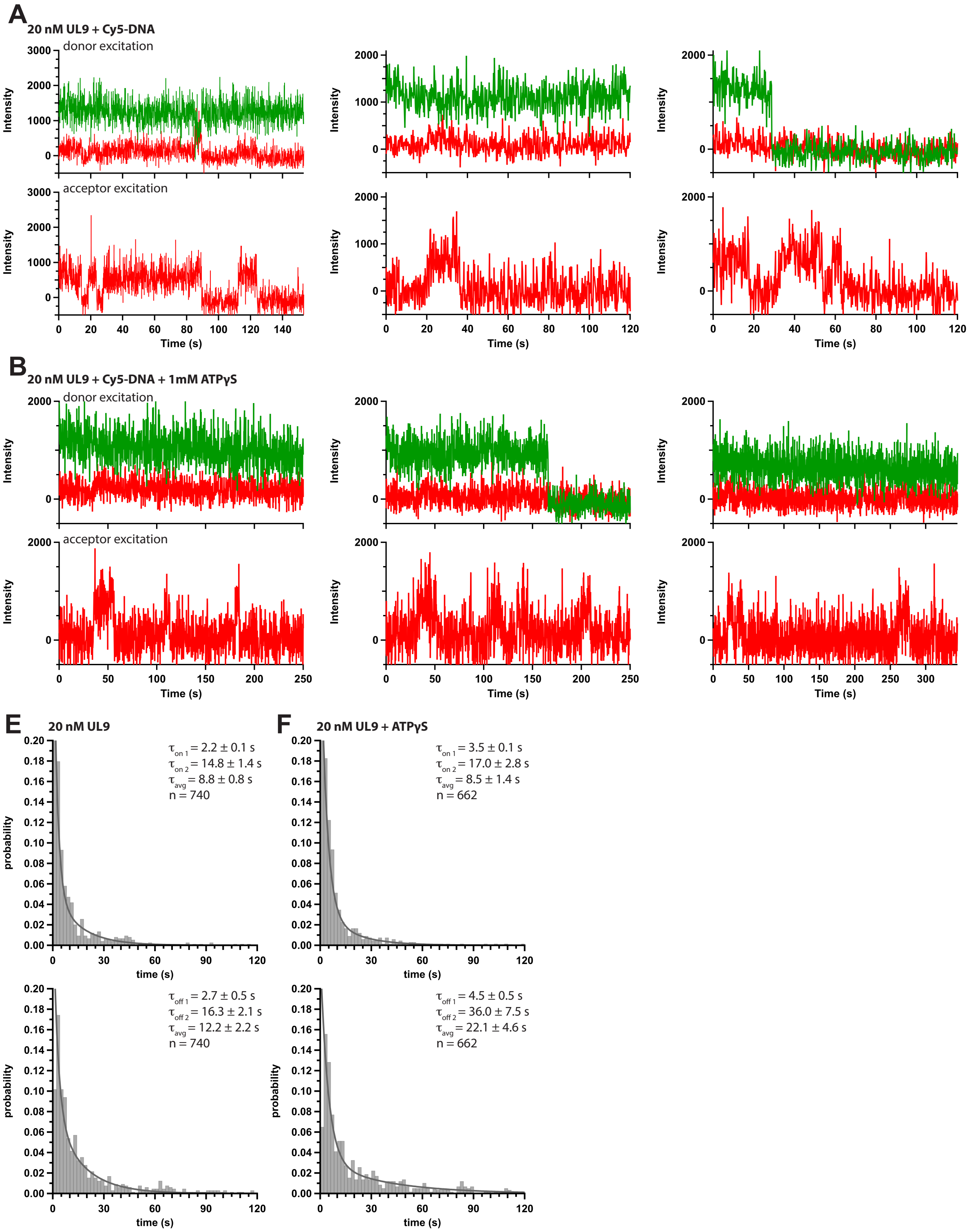


**Figure S5: UL9 captures DNA in trans.** Additional smFRET traces under donor (top) or acceptor excitation in the presence of 20 nM UL9, 10 nM Cy5-DNA in the absence (**A**) or presence (**B**) of 1 mM ATPγS. Dwell time analysis of Cy5-DNA in the absence (**E**) or presence (**F**) of 1 mM ATPγS.
